# Context-dependent Foxa2 activity maintains floor plate fate and tunes Sonic hedgehog signaling to regulate neural progenitor differentiation

**DOI:** 10.64898/2026.06.21.733624

**Authors:** Aarti Kejriwal, Mindy Kim, Lucas D. Lo Vercio, Peng Huang

## Abstract

The developing spinal cord contains both neural and non-neural tissues that arise within a shared morphogen signaling environment, raising the question of how distinct lineage identities are established and maintained. A striking example of this is the floor plate (FP), a mesoderm-derived, non-neural midline structure that functions as a critical signaling center adjacent to neural progenitor domains. Here, using zebrafish, we show that the pioneer transcription factor *foxa2* is expressed in the FP, adjacent p3 neural progenitors, and p3-derived Kolmer-Agduhr” (KA”) interneurons. Loss of *foxa2* results in a complete loss of canonical FP identity and an expansion of p3 progenitors. Lineage tracing reveals that, in the absence of *foxa2*, FP cells undergo a fate transformation into neuron-producing p3-like cells, indicating that Foxa2 functions as a lineage barrier to preserve non-neural FP identity. In contrast, within the neural lineage, loss of *foxa2* leads to elevated Sonic hedgehog (Shh) pathway activity and impaired KA” differentiation, suggesting that Foxa2 negatively regulates Shh responsiveness. Conversely, Foxa2 overexpression induces ectopic FP and KA” marker expression in a stage-dependent manner. Together, our findings reveal a dual role for Foxa2 in maintaining the non-neural FP lineage while fine-tuning morphogen responsiveness in neighboring neural progenitors during spinal cord development.

## INTRODUCTION

The vertebrate spinal cord is patterned along the dorsoventral axis by opposing gradients of morphogens that influence the fate and identity of neural progenitor cells^1–4^. Bone morphogenetic proteins (BMPs) and Wnts are secreted from the dorsal roof plate (RP), while Sonic hedgehog (Shh) is secreted from ventral sources, including the notochord and the floor plate (FP)^2–7^. The timing and levels of these opposing morphogen gradients induce different combinations of transcription factors, which in turn define distinct progenitor domains^1–3,6,8,9^. These neural progenitors give rise to different interneurons and motor neuron populations^10^. Although the spinal cord primarily consists of neural tissue, specialized midline structures such as the RP and FP adopt non-neural identities. Rather than differentiating into neurons, these cells function as critical signaling centers that influence the fate and organization of surrounding neural progenitors^11,12^. This raises a fundamental question: how are these non-neural tissues specified and maintained in a neurogenic environment?

In zebrafish, the FP, also known as the medial floor plate (MFP), together with the notochord and hypocord, originates from the axial mesodermal progenitors^13,14^. Lineage tracing studies have demonstrated that these progenitor cells arise from the shield during gastrulation, a region analogous to the node organizer in mice^13–15^. Adjacent to the FP lies the lateral floor plate (LFP), which derives from the neuroectoderm and, in a sequential manner, gives rise to Kolmer-Agduhr” (KA”) interneurons, V3 interneurons, and intraspinal serotonergic neurons^16–20^. Functionally and structurally, the LFP in zebrafish corresponds to the p3 progenitor domain described in other vertebrate species^16,20–22^. For consistency of nomenclature across species, we refer to the MFP as the FP and the LFP as the p3 progenitor domain throughout this manuscript.

Although FP and p3 cells arise from distinct embryonic lineages and follow distinct developmental trajectories, both domains share expression of the pioneer transcription factor Foxa2 across vertebrate species^21,23–27^. Foxa2 is a highly conserved forkhead box transcription factor with roles in multiple tissues during embryonic development^28–30^. Genetic studies in mouse have shown that Foxa2 is essential for the formation of the node and midline structures, including the notochord and floor plate^31–34^. Foxa2 mutant mice embryos do not form a distinct node, lack a notochord, and eventually die by embryonic day (E) 10-11^31–33^. Beyond its roles in midline development, Foxa2 is also critical for endoderm development, where its early enhancer binding establishes competence for liver and pancreas specification^29,35–40^.

As a member of the Foxa pioneer transcription factor family, Foxa2 can displace histones and render chromatin accessible to lineage-specific regulators^30,39,41^. However, its ability to stably open chromatin often requires cooperation with additional factors, suggesting that its function is tissue- and context-dependent^42^. This raises the possibility that Foxa2 may perform distinct regulatory roles in different cellular contexts within the ventral spinal cord, including the FP and the adjacent p3 domain, structures that arise from separate embryonic lineages yet share Foxa2 expression. While Foxa2’s role in the induction of midline structures, including the FP, has been well studied^24,33,43–45^, its precise functions in the spinal cord remain poorly understood. Here, we address this gap by investigating the distinct roles of Foxa2 in both the FP and the p3 domain of the developing spinal cord.

Using zebrafish as a model, we show that *foxa2* is essential for specifying FP identity and preventing lineage switching of FP cells into neuron-producing cells. Loss of *foxa2* results in FP cells adopting neuronal fates, while gain-of-function induces ectopic expression of FP markers. Furthermore, we uncover a distinct role for Foxa2 in the p3 domain, where it modulates Shh signaling responsiveness and influences neuronal differentiation. Together, these findings reveal that Foxa2 acts as a key regulator of ventral spinal cord patterning, maintaining FP identity while fine-tuning neurogenic programs in adjacent neural progenitor domains.

## RESULTS

### *foxa2* is expressed in both FP and p3 domains in the zebrafish spinal cord

In zebrafish, the FP is a single row of cells flanked by two adjacent rows forming the p3 domain, which gives rise to KA” and V3 interneurons (Fig. 1A-B). Double fluorescent in situ hybridization and immunohistochemistry revealed that *foxa2* was expressed in the FP, marked by *shha* expression, as well as in the *nkx2.2a*⁺ p3 domain (Fig. 1C-D). Additionally, *foxa2* was detected in *gata2:GFP*-positive KA” interneurons but was absent from *vglut2a:GFP*-positive V3 interneurons (Fig. 1E). Interestingly, KA” cells exhibited higher *foxa2* expression compared to neighboring p3 progenitors, as revealed by both fluorescent in situ hybridization and immunohistochemistry (Fig. 1E), suggesting that *foxa2* is upregulated during KA” differentiation. Notably, while *foxa2* transcripts were detected in p3 progenitors by fluorescent in situ hybridization, Foxa2 protein was detectable only in FP cells and KA” interneurons by immunohistochemistry, with little protein signal observed in p3 progenitors at 30 hours post fertilization (hpf) (Fig. 1E). Together, our expression analysis indicates that *foxa2* functions in both neural and non-neural cells of the spinal cord.

**Figure 1.**
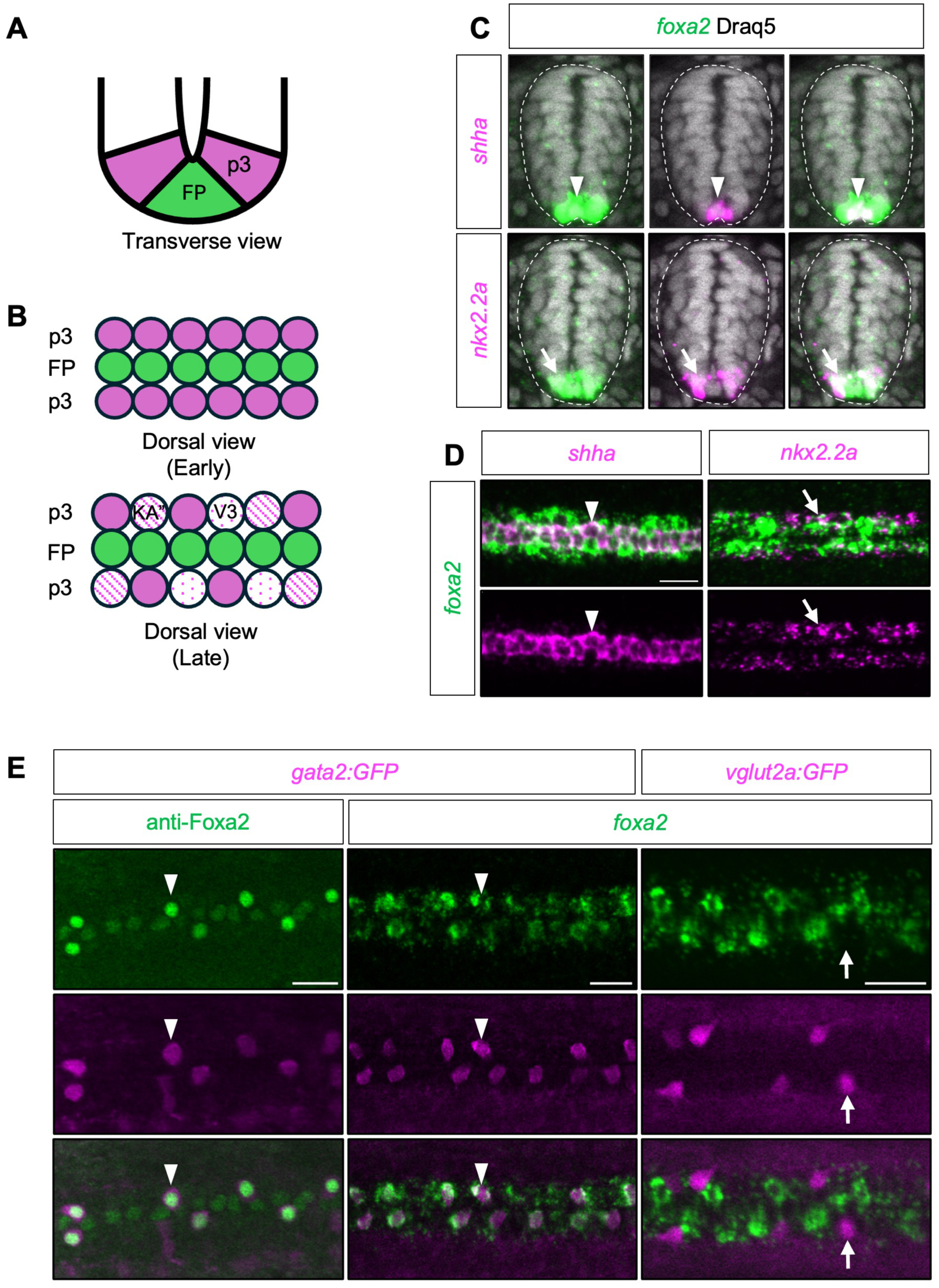
Expression analysis of *foxa2* in the zebrafish spinal cord. **(A)** Schematic transverse view of the zebrafish spinal cord illustrating the organization of the FP and p3 domains at 24 hpf. **(B)** Schematic dorsal views of the spinal cord at early and late developmental stages. At early stages between 12 to 18 hpf, the FP occupies the ventral midline and is flanked by p3 progenitors. At later stages between 24 and 48 hpf, differentiated neuronal populations, including KA” and V3 interneurons, emerge lateral to the FP. **(C)** Transverse sections showing fluorescent in situ hybridization for *foxa2* (green) together with *shha* or *nkx2.2a* (magenta), with Draq5 marking nuclei (grey), revealing *foxa2* expression in the FP (arrowheads) and the adjacent p3 domain (arrows). Dashed outlines indicate the spinal cord. *n* = 15 per staining. **(D)** Dorsal views of the spinal cord with co-labeling of *foxa2* (green) with *shha* or *nkx2.2a* (magenta), showing *foxa2* expression in both FP (arrowheads) and p3 (arrows) domains. *n* = 15 per staining. **(E)** Dorsal views of the spinal cord from *gata2:GFP* (magenta) and *vglut2a:GFP* (magenta) transgenic embryos showing Foxa2 protein (left column; green) and *foxa2* mRNA (middle and right columns; green) expression. *foxa2* is expressed in KA” interneurons (arrowheads) but not in V3 interneurons (arrows). *n* ≥ 10 per staining. All images were acquired from embryos fixed at 30 hpf. Scale bars: 20 μm.

### Absence of Foxa2 leads to FP loss and p3 expansion

To investigate the role of *foxa2* in FP development, we used two loss-of-function approaches: a *foxa2* mutant (*foxa2^-/-^*) carrying a single C-to-A point mutation that introduces a premature stop codon, resulting in a truncated forkhead box DNA-binding domain^44^; and a morpholino-mediated knockdown of *foxa2* (*foxa2^MO^*). Foxa2 immunostaining confirmed complete loss of Foxa2 protein in both *foxa2^-/-^* mutants and *foxa2^MO^* embryos (Fig. S1A). In situ hybridization revealed a complete loss of FP markers *shha*, *shhb*, and *spon1b* in *foxa2^-/-^* mutants (Fig. 2A). Consistently, in embryos subjected to morpholino-mediated knockdown of *foxa2* (*foxa2^MO^*), *shhb* expression was entirely absent, while *shha* and *spon1b* expressions were markedly reduced (Fig. S1B). Live imaging using the transgenic line *ccn2a:EGFP*, which labels both the notochord and the floor plate at 30 hpf, revealed a complete absence of EGFP expression in the presumptive FP region in *foxa2^MO^* embryos (Fig. 2B). Importantly, EGFP expression in the notochord remained unaffected (Fig. 2B), indicating that the *foxa2* knockdown specifically disrupts FP expression without impacting notochord labeling. Together, these findings demonstrate that *foxa2* is essential for FP development.

**Figure 2.**
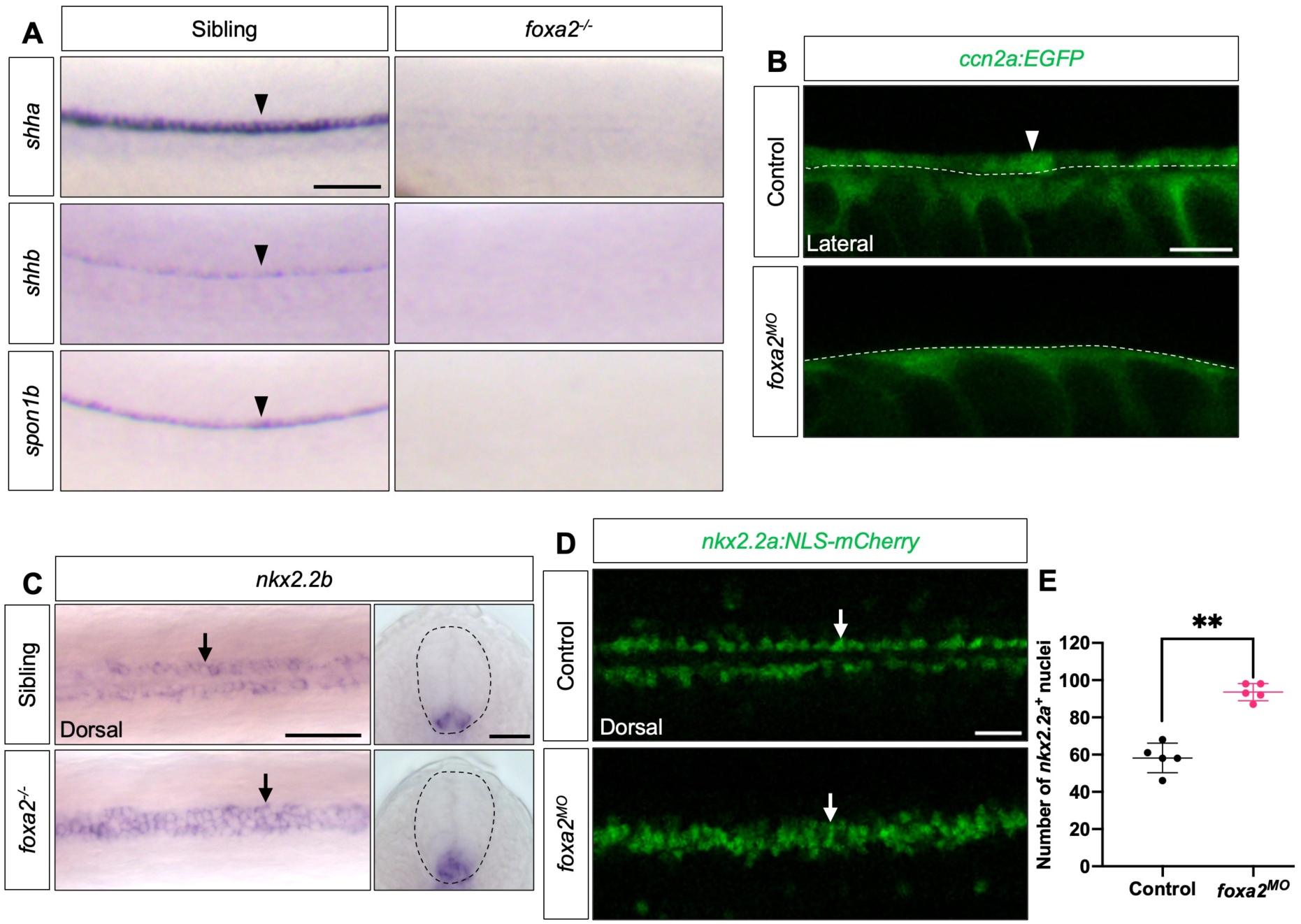
Loss of Foxa2 leads to loss of FP identity and expansion of the p3 domain. **(A)** Whole-mount in situ hybridization for the FP markers *shha*, *shhb*, and *spon1b* in sibling (*foxa2^+/+^* or *foxa2^+/-^*) and *foxa2^-/-^* mutant embryos, showing a complete loss of FP marker expression (arrowheads) in the absence of Foxa2. *n* = 25 per staining. **(B)** Live imaging of transgenic embryos expressing *ccn2a:EGFP*, which labels the floor plate and notochord (lateral views), in control and *foxa2* morpholino-injected (*foxa2^MO^*) embryos. Dashed lines mark the upper boundary of the notochord. *ccn2a:EGFP* expression is absent from the FP (arrowheads) in *foxa2^MO^* embryos. *n* = 10 per condition. **(C)** Whole-mount in situ hybridization for the p3 marker *nkx2.2b* in sibling controls and *foxa2^-/-^*mutants. Dorsal and transverse views show expansion of *nkx2.2b* expression (arrows) into the ventral midline in *foxa2^-/-^* embryos. Dashed outlines indicate the spinal cord. *n* = 20 per staining. **(D)** Live imaging of transgenic embryos expressing *nkx2.2a:NLS-mCherry*, which labels p3 progenitors (dorsal views), in control and *foxa2^MO^* embryos. mCherry^+^ cells are expanded in *foxa2^MO^* embryos. *n* = 30 per condition. **(E)** Quantification of *nkx2.2a^+^* nuclei in control and *foxa2^MO^* embryos following *nkx2.2a* in situ hybridization and Draq5 nuclear staining. *nkx2.2a^+^* nuclei were counted within a defined three-somite region of the spinal cord in fixed embryos. Each data point represents one embryo. *n* = 5 embryos per condition. Data are plotted as mean ± SD. Statistics: Mann-Whitney U test. Significance: p < 0.01 (**). All images were acquired from embryos fixed or imaged at 30 hpf. Scale bars: 100 μm (A); 20 μm (B, D); 50 μm (dorsal view) and 20 μm (transverse view) (C).

To assess the impact of Foxa2 loss on the p3 progenitor domain, we performed in situ hybridization using p3-specific markers in *foxa2^-/-^* mutants compared with their sibling controls (*foxa2^+/+^* and *foxa2^+/-^*), as well as in *foxa2^MO^* embryos compared with uninjected sibling controls (Fig. 2C and Fig. S1C). Interestingly, the p3 marker *nkx2.2b* was expanded in Foxa2-deficient embryos (Fig. 2C and Fig. S1C). Transverse sections of *nkx2.2b*-stained embryos revealed that, in the absence of Foxa2, p3 markers were expressed in the region normally occupied by FP cells (Fig. 2C and Fig. S1C). Live imaging using the *nkx2.2a:NLS-mCherry* transgenic line showed an increased number of mCherry^+^ cells, with three rows of labeled cells in *foxa2^MO^* compared to two in controls (Fig. 2D). Quantification of embryos co-labeled with *nkx2.2a* and Draq5 nuclear staining revealed a 61% increase in the number of p3 cells in *foxa2^MO^*compared to controls (Fig. 2E). Together, these results indicate that Foxa2 is required for proper floor plate development, and that loss of Foxa2 leads to an expansion of the p3 progenitor domain.

### Lineage tracing reveals a FP-to-p3 fate switch in the absence of Foxa2

The expansion of the p3 domain in the absence of Foxa2 suggests two possibilities: (1) FP precursor cells fail to reach their correct midline location, allowing neighboring p3 cells to expand into the vacant territory; or (2) FP cells reach the midline but fail to express canonical FP markers and instead adopt a p3-like fate. To distinguish between these scenarios, we performed fate-mapping experiments. Previous studies have shown that FP cells originate from the embryonic shield, along with other axial mesodermal tissues such as the notochord and hypochord^13–15,45,46^. To trace the fate of shield-derived cells, we used two independent lineage-labeling strategies. First, we injected caged fluorescein dextran at the one-cell stage and photoactivated a small group of shield cells at 6 hpf using a 405-nm laser (Fig. 3A). Second, we injected Kaede mRNA at the one-cell stage and photoconverted shield cells from green (Kaede^green^) to red (Kaede^red^) fluorescence at 6 hpf. In control embryos imaged at 24 hpf, fluorescein- or Kaede^red^-labeled cells contributed to both the notochord and floor plate, as expected (Fig. 3B). In both *foxa2^MO^*and *foxa2^-/-^* embryos, shield-derived cells still localized to the midline above the notochord (Fig. 3B, 3C, and 3E). This indicates that these cells reach the correct position but fail to differentiate into FP cells, supporting a fate switch rather than a migration defect.

**Figure 3.**
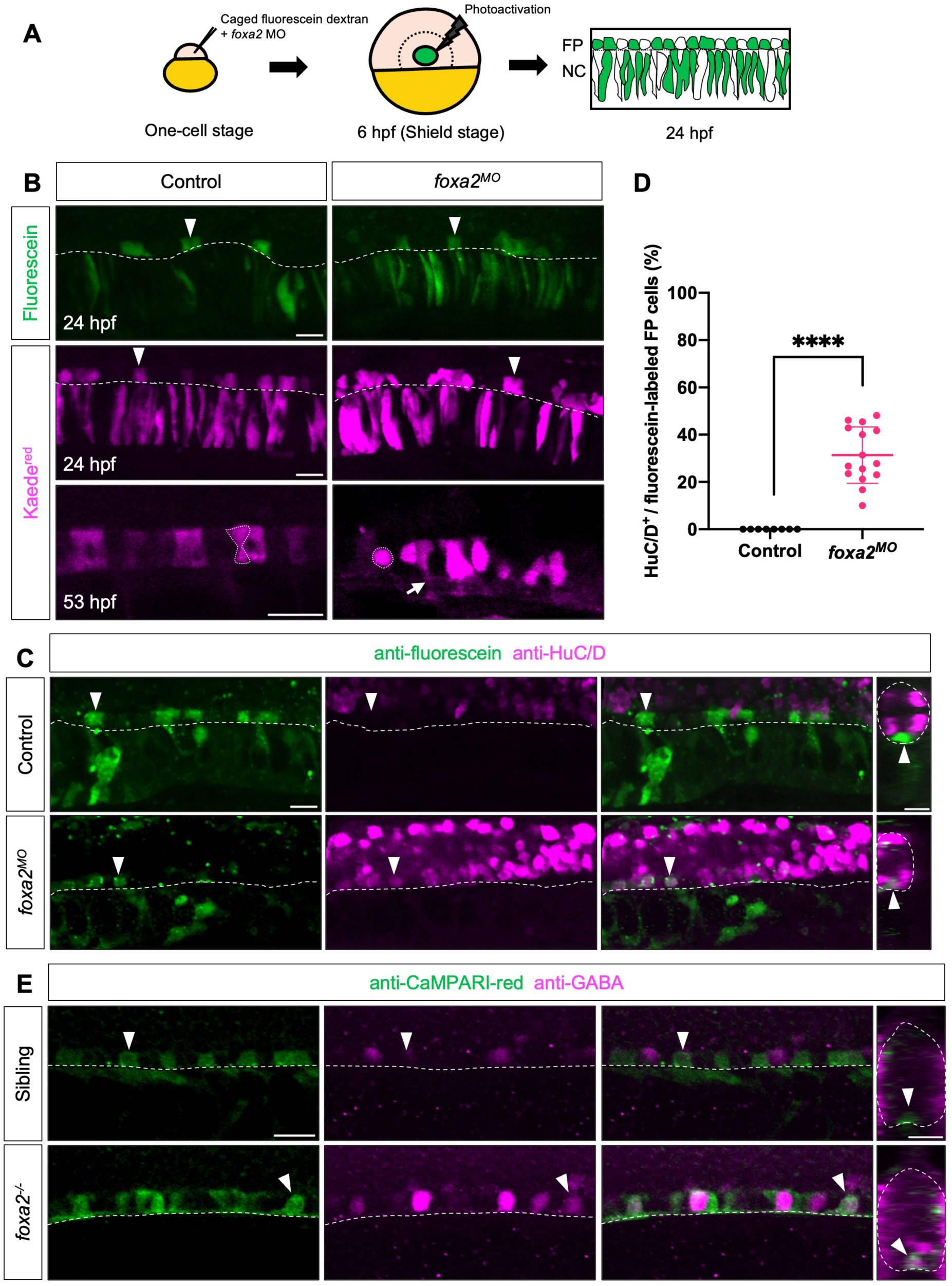
Lineage tracing reveals fate switching of FP cells in the absence of Foxa2. **(A)** Schematic of the lineage-tracing strategy. Embryos were injected at the one-cell stage with caged fluorescein dextran with or without *foxa2* morpholino (*foxa2^MO^*). Photoactivation was performed at the shield stage (6 hpf) to label the central region of the shield, resulting in mosaic labeling of FP and notochord (NC) cells at 24 hpf. (**B)** Representative images of lineage-traced FP cells (arrowheads) in control and *foxa2^MO^* embryos at 24 hpf using fluorescein photoactivation (green) and Kaede photoconversion (Kaede^red^, magenta). In control embryos, labeled FP cells exhibit the characteristic hourglass-shaped morphology (dotted outline) at 53 hpf, whereas in *foxa2^MO^*embryos, labeled cells adopt neuron-like morphologies (dotted outline) with axon-like extensions (arrow). *n* = 20 (fluorescein), 20 (Kaede, 24 hpf), and 5 (Kaede, 53 hpf) embryos. **(C)** Immunofluorescence staining for fluorescein (green) and the pan-neuronal marker HuC/D (magenta) in control and *foxa2^MO^*embryos at 30 hpf, showing neural differentiation of fluorescein-labeled cells (arrowheads) following Foxa2 loss. *n* = 8 (control) and 14 (*foxa2^MO^*) embryos. **(D)** Quantification of the percentage of HuC/D⁺ cells among fluorescein-labeled cells in control sibling and *foxa2^MO^* embryos. Each data point represents one embryo. *n* = 8 (control) and 14 (*foxa2^MO^*) embryos. Data are plotted as mean ± SD. Statistics: Mann–Whitney U test. Significance: p < 0.0001 (****). **(E)** Immunofluorescence staining for Kaede^red^ using anti-CaMPARI-red (green) and KA” interneuron marker anti-GABA (magenta) in control sibling and *foxa2^-/-^* embryos at 30 hpf, showing acquisition of GABAergic neuronal identity by Kaede^red^ lineage-traced cells (arrowheads) in the absence of Foxa2. *n* = 3 (sibling) and 4 (*foxa2^-/-^*) embryos. Dashed lines mark the boundary between the FP and notochord in lateral views and outline the spinal cord in transverse views. Scale bars: 20 μm.

To test whether these prospective FP cells lacking Foxa2 adopt a p3 fate and contribute to neurons such as KA” and p3 interneurons, we first examined cell morphology at 53 hpf. In controls, Kaede^red^-labeled cells above the notochord showed the characteristic hourglass-like FP morphology^47^ (Fig. 3B). In contrast, labeled cells in *foxa2^MO^* embryos lacked this morphology and instead exhibited neuronal-like shapes, including axonal projections (Fig. 3B). To determine if these shield-derived cells generate neurons, we performed double labeling with the pan-neuronal marker anti-HuC/D and an anti-fluorescein antibody (Fig. 3C). Quantification showed that ∼31% of fluorescein-labeled cells in the presumptive FP region in *foxa2^MO^* embryos were also HuC/D-positive, while no double-positive cells were observed in controls (Fig. 3C-D). These results suggest that, in the absence of Foxa2, prospective FP cells adopt a p3-like fate and contribute to neuronal lineages. To further support this observation, we used the Kaede-based fate-mapping approach followed by immunostaining with anti-GABA to label KA” interneurons and anti-CaMPARI-red to detect Kaede^red^-labeled cells. In *foxa2*-deficient embryos, but not in controls, we observed co-localization of GABA and Kaede^red^ (Fig. 3E), indicating that shield-derived cells can adopt a KA” neuronal fate in the absence of Foxa2. Together, these findings support a fate-switching model in which Foxa2 is essential for specifying FP identity and repressing neuronal program of midline progenitors.

### Fate-switched FP cells in the absence of Foxa2 respond to different levels of Shh signaling

In zebrafish, FP specification occurs independently of Shh signaling^21,48^, even though the FP later serves as a major source of Shh ligands. In contrast, Shh signaling is essential for patterning the surrounding ventral neural progenitor domains, including the p3 domain^16^. Since the FP transforms into a p3-like domain in the absence of Foxa2, we asked whether these transformed cells gain responsiveness to Shh. Using the Shh reporter line *ptc2:Kaede*^49,50^, which marks cells responding to Shh, we compared Shh signaling activity in wild-type and *foxa2^MO^*embryos. In controls, Kaede expression was absent from the ventral-most spinal cord corresponding the FP region (Fig. 4A), consistent with FP cells being Shh-insensitive. In contrast, *foxa2^MO^* embryos exhibited Kaede expression in these ventral cells (Fig. 4A), suggesting that the transformed FP cells now respond to Shh signaling.

**Figure 4.**
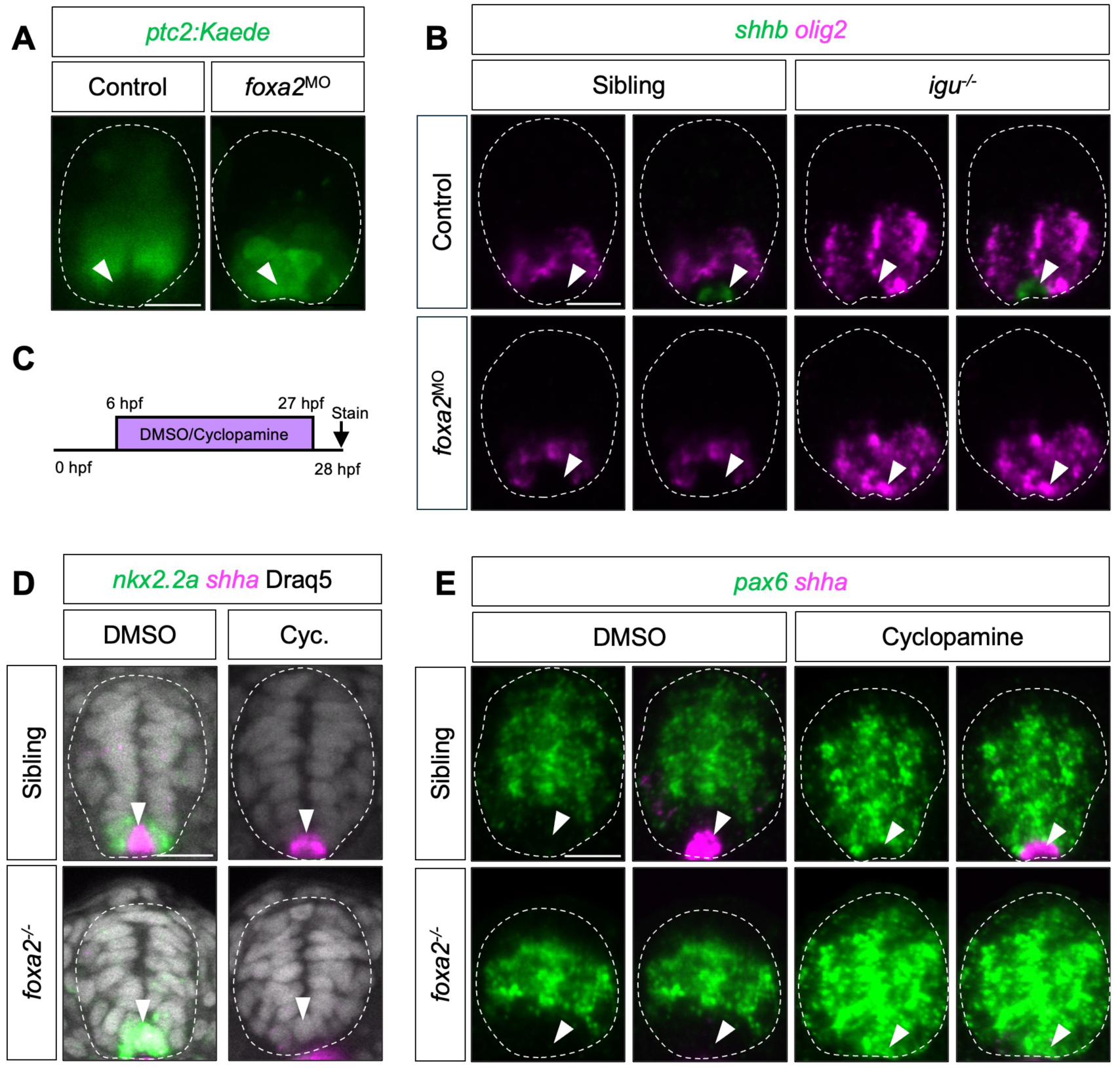
Fate-switched FP cells acquire Shh signaling responsiveness in the absence of Foxa2. **(A)** Representative transverse sections showing *ptc2:Kaede* reporter expression in control and *foxa2^MO^* embryos at 28 hpf, revealing expanded reporter expression at the ventral midline (arrowheads) in *foxa2^MO^*embryos. *n* = 12 embryos per condition. **(B)** Fluorescent in situ hybridization for *shhb* (green) and *olig2* (magenta) in uninjected and *foxa2^MO^*-injected sibling controls (*igu^+/+^* or *igu^+/-^*) and homozygous *igu^-/-^* mutants at 30 hpf. *n* ≥ 10 per staining. **(C)** Schematic of cyclopamine treatment regimen. Embryos were treated with DMSO or cyclopamine from 6 hpf to 27 hpf, followed by fixation and staining at 28 hpf. **(D)** Fluorescent in situ hybridization for *nkx2.2a* (green) and *shha* (magenta) in sibling controls (*foxa2^+/+^* or *foxa2^+/-^*) and *foxa2^-/-^*mutants treated with DMSO or cyclopamine (Cyc.). Nuclei are labeled by Draq5 staining. *n* ≥ 5 per staining. **(E)** Fluorescent in situ hybridization for the dorsal spinal cord marker *pax6* (green) and *shha* (magenta) in control and *foxa2^-/-^* mutant embryos treated with DMSO or cyclopamine. *n* ≥ 5 per staining. In (A, B, D, E), transverse views are shown with dashed outlines indicating the spinal cord, and arrowheads point to the ventral midline of the spinal cord. Scale bars: 20 μm.

Based on the gain of Shh responsiveness in the absence of Foxa2, we predict that transformed FP cells could respond to different levels of Shh signaling and adopt distinct neural fates. To test this, we manipulated Shh signaling genetically and pharmacologically. We first used zebrafish *iguana* mutants (*igu^-/-^*), in which *iguana* encodes the ciliary basal body protein Dzip1^51–56^. In zebrafish, loss of primary cilia, as in *iguana* mutants, results in a reduction of high-level Shh response accompanied by an expansion of low-level Shh pathway activity^52,57^. This expanded Shh signaling is insufficient to induce high-threshold, Shh-dependent domains such as p3, but is sufficient to allow expansion of the motor neuron progenitor (pMN) domain^50,52^. Consistent with this, *igu^-/-^*embryos exhibited expansion of the *olig2^+^* pMN domain and failed to form the *nkx2.2a^+^* p3 domain, while FP cells retained normal *shhb* expression (Fig. 4B and Fig. S2A). Strikingly, when *foxa2* was knocked down in *igu^-/-^* embryos (*igu^-/-^* + *foxa2^MO^*), prospective FP cells lost *shhb* expression and instead ectopically expressed pMN marker *olig2* in the ventral-most spinal cord, suggesting a fate switch to pMN identity (Fig. 4B). These results suggest that, in the absence of Foxa2, FP cells adopt a p3-like fate under higher Shh signaling (*foxa2^MO^*alone) but switch to a pMN fate under lower Shh signaling (*igu^-/-^*+ *foxa2^MO^*).

To complement these findings, we used cyclopamine, a specific Smoothened inhibitor, to completely block Shh signaling. Embryos from *foxa2^+/-^*intercrosses were treated with DMSO or cyclopamine from 6 to 27 hpf and fixed at 28 hpf (Fig. 4C). As expected, cyclopamine treatment eliminated the p3 marker *nkx2.2a* expression in the ventral spinal cord (Fig. 4D). In *foxa2^+/+^*and *foxa2^+/-^* control embryos treated with cyclopamine, FP cells retained *shha* expression despite dorsalization of the neighboring ventral cells, as indicated by the expanded *pax6* (Fig. 4E) and *irx3* expression (Fig. S2B). In contrast, in cyclopamine-treated *foxa2^-/-^* embryos, prospective FP cells in the ventral-most spinal cord no longer expressed *shha* and instead expressed the dorsal markers *pax6* and *irx3*, indicating a transformation into dorsal progenitor identity (Fig. 4E and Fig. S2B).

Collectively, these experiments demonstrate that Foxa2 prevents FP cells from switching to neural progenitor fates and from responding to Shh signaling. In its absence, FP cells become Shh responsive, and their fate is determined by the level of Shh signaling activity: they adopt p3 identity under high Shh, pMN identity under intermediate Shh, and dorsal identities when Shh signaling is absent.

### Foxa2 is required for proper differentiation of p3-derived neurons

Although loss of Foxa2 results in an expansion of the *nkx2.2a/b^+^*p3 progenitor domain, it is unclear whether these expanded progenitors differentiate normally. This distinction is important because p3 progenitors must switch off Shh responsiveness before differentiating into KA” and V3 interneurons^16,18,50^. We therefore next examined whether loss of Foxa2 alters Shh pathway responsiveness and the production of p3-derived neurons.

We first assessed Shh signaling activity in the spinal cord using the *ptc2:Kaede* reporter line, comparing *foxa2^MO^* embryos with controls (Fig. 5A). To quantify *ptc2:Kaede* expression, we adapted a modified MATLAB-based analysis pipeline derived from Cimtool^58,59^. This approach extracted fluorescence intensity profiles across the ventral-dorsal axis of the segmented spinal cord, enabling quantitative comparison of Shh signaling responses between conditions. Using this method, we detected significantly elevated *ptc2:Kaede* levels in the ventral-most spinal cord of *foxa2^MO^*embryos compared to controls (Fig. 5B-C), suggesting that Foxa2 negatively regulates Shh response. Notably, this increase occurred despite the loss of the FP, one of the two primary sources of Shh ligands, in *foxa2^MO^* morphants. In addition to the overall increase in reporter activity, *foxa2^MO^* embryos exhibited a significantly steeper ventral-dorsal *ptc2:Kaede* gradient, as indicated by a larger negative slope compared to controls (Fig. 5D). This pattern suggests a ventrally concentrated Shh response that declines sharply toward dorsal regions, rather than forming a broader gradient like in wild-type embryos.

**Figure 5.**
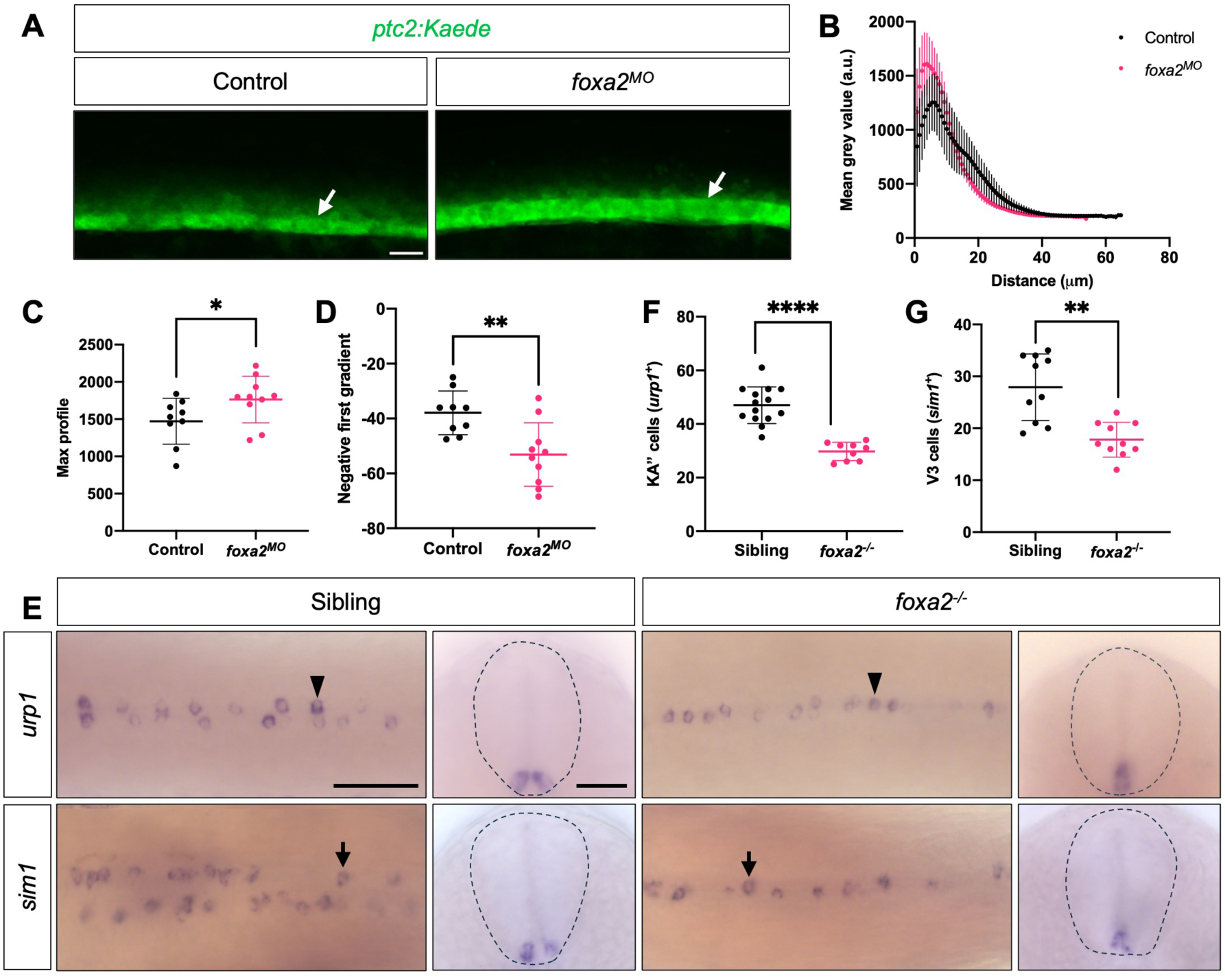
Foxa2 loss increases Shh signaling response and disrupts p3-derived interneuron differentiation. **(A)** Representative lateral views of live *ptc2:Kaede* reporter expression in control and *foxa2^MO^*embryos at 28 hpf, showing increased Shh pathway activity (arrowheads) along the ventral spinal cord following Foxa2 loss. Quantifications of *ptc2:Kaede* expression are shown in (B-D). *n* = 9 (control) and 10 (*foxa2^MO^*) embryos. **(B)** Averaged *ptc2:Kaede* fluorescence intensity profiles along the dorsoventral axis of the spinal cord in control and *foxa2^MO^* embryos. Fluorescence intensity was quantified along the segmented spinal cord and plotted as mean ± SD as a function of distance from the ventral midline. **(C)** Quantification of the maximum *ptc2:Kaede* fluorescence intensity in control and *foxa2^MO^* embryos. Each data point represents the median of multiple measurements from a single embryo. **(D)** Quantification of the slope of the ventral-to-dorsal *ptc2:Kaede* fluorescence gradient in control and *foxa2^MO^* embryos, showing a steeper negative gradient upon Foxa2 loss. Each data point represents the median of multiple measurements from a single embryo. **(E)** Whole-mount in situ hybridization for the KA” interneuron marker *urp1* and the V3 interneuron marker *sim1* in sibling controls (*foxa2^+/+^*or *foxa2^+/-^*) and *foxa2^-/-^* mutant embryos at 30 hpf, revealing a reduced number of KA” (arrowheads) and V3 (arrows) cells in the absence of Foxa2. Dorsal and transverse views are shown, with dashed lines outlining the spinal cord. *n* = 25 per staining. **(F)** Quantification of KA” interneuron numbers in sibling controls and *foxa2^-/-^* mutants at 30 hpf. Each data point represents one embryo. *n* = 14 (control) and 9 (*foxa2^-/-^*) embryos. **(G)** Quantification of V3 interneuron numbers in sibling controls and *foxa2^-/-^* mutants at 30 hpf. Each data point represents one embryo. *n* = 10 embryos per condition. Data in (C, D, F, G) are plotted as mean ± SD. Statistics: Mann–Whitney U test. Significance: p < 0.05 (*), p < 0.01 (**), p < 0.0001 (****). Scale bars: 20 μm (A); 50 μm (dorsal view) and 20 μm (transverse view) (E).

Our previous studies have shown that spinal cord progenitors must attenuate their Shh signaling response in order to differentiate into neurons^16,50^. Given the elevated Shh response observed in Foxa2-deficient embryos, we hypothesized that this may impair the differentiation of p3-derived interneurons. To test this, we performed in situ hybridization for the KA” and V3 interneuron markers, *urp1* and *sim1*, respectively, in both *foxa2^MO^* and *foxa2^-/-^*embryos (Fig. 5E and Fig. S3A). In the absence of Foxa2, the number of both KA” and V3 interneurons was significantly reduced (Fig. 5F-G and Fig. S3B-C). Interestingly, the neurons that did form were positioned along the midline rather than in the two lateral rows characteristic of wild-type embryos (Fig. 5E and Fig. S3A). This is particularly striking because, despite the increase in p3 progenitors observed in the absence of Foxa2, the number of differentiated interneurons was reduced. These findings suggest that elevated Shh responsiveness in the absence of Foxa2 compromises the differentiation of p3-derived neurons.

### Ectopic Foxa2 expression is sufficient to induce FP and KA” fate markers

To complement the loss-of-function analyses, we performed gain-of-function experiments using a heat-shock inducible *hsp:foxa2-P2A-EGFP* construct (*foxa2^GOF^*). Embryos were injected at the one-cell stage, heat-shocked at 26 hpf, and fixed 6 hours later. Immunohistochemistry confirmed robust ectopic Foxa2 expression that colocalized with EGFP, indicating successful overexpression (Fig. 6A). Heat-shock-induced Foxa2 overexpression at 10 hpf resulted in ectopic expression of multiple FP markers, including *shha*, *shhb*, and *spon1b*, in EGFP-positive *foxa2^GOF^*cells at 16 hpf (Fig. 6B), demonstrating cell-autonomous mode of action. Strikingly, Foxa2 overexpression also triggered widespread ectopic expression of the KA” interneuron markers *urp1* and *urp2* throughout the embryo (Fig. 6B and Fig. S4A-B). To quantify the induction of different identity markers following Foxa2 overexpression, we calculated the proportion of EGFP-positive cells expressing each marker (Fig. 6C). Nearly all EGFP-positive cells (∼95%) expressed *urp1*, whereas only a smaller fraction (∼50%) expressed FP markers (*shha*, *shhb*, and *spon1b*) (Fig. 6C). To determine whether this difference reflects temporal constraints on FP specification, we performed a stage-specific induction of Foxa2. Foxa2 overexpression at early stages (11 hpf) strongly induced ectopic *shha* expression, while induction at mid stages (24 hpf) produced a markedly reduced response, and late induction (48 hpf) did not induce floor plate marker expression (Fig. 6D-E). In contrast, *urp1* was consistently upregulated at all stages examined (Fig. 6D and 6F), indicating that Foxa2 overexpression robustly promotes KA” interneuron identity across developmental time, while FP marker induction is temporally restricted.

**Figure 6.**
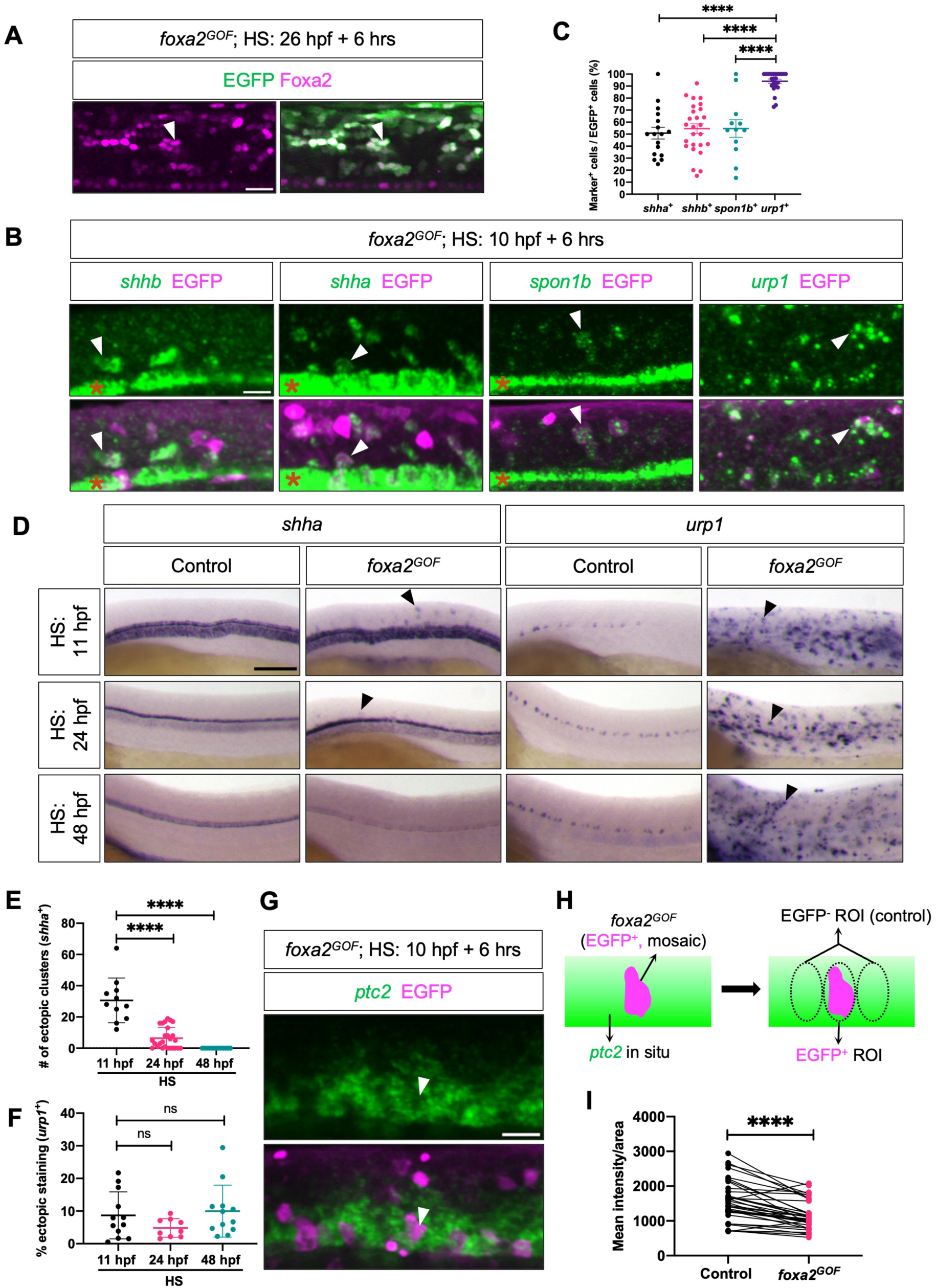
Foxa2 overexpression induces FP and KA” programs while attenuating Shh signaling. **(A)** Representative images of embryos injected with *hsp:foxa2-P2A-EGFP* (referred as *foxa2^GOF^*) following heat shock at 26 hpf, showing co-staining for EGFP (green) and Foxa2 (magenta) (arrowheads) at 32 hpf. *n* = 12 embryos. **(B)** Fluorescent in situ hybridization for the FP markers *shhb*, *shha*, and *spon1b*, and the KA” interneuron marker *urp1* (green), together with immunofluorescence staining for EGFP (magenta), in *foxa2^GOF^* embryos following heat shock at 10 hpf and fixation at 16 hpf. Asterisks indicate endogenous expression domains in the FP, which were excluded from ectopic expression scoring. Arrowheads indicate ectopic marker expression associated with EGFP^+^ Foxa2-overexpressing cells. *n* ≥ 12 per staining. **(C)** Quantification of the percentage of EGFP⁺ cells co-expressing the FP and KA” markers in the experiments shown in (B). Each data point represents one embryo. *n* = 12-26 embryos per marker. **(D)** Whole-mount in situ hybridization for *shha* and *urp1* in control and *foxa2^GOF^*embryos following heat shock at the indicated developmental stages (11, 24, and 48 hpf) and fixation 6 hours later. Ectopic FP and KA” marker expression following Foxa2 overexpression is indicated by arrowheads. *n* ≥ 10 (*shha*) and 12 (*urp1*) per staining. **(E)** Quantification of the number of ectopic *shha^+^* clusters in the experiment shown in (D). Each data point represents one embryo. *n* = 11-26 embryos per condition. **(F)** Quantification of the percentage area of each embryo covered by ectopic *urp1* staining in the experiment shown in (D). Each data point represents one embryo. *n* = 9-12 embryos per condition. **(G)** Fluorescent in situ hybridization for the Shh signaling target *ptc2* (green) together with EGFP (magenta; arrowheads) in *foxa2^GOF^* embryos following heat shock at 10 hpf and fixation at 16 hpf. Quantification of this experiment is shown in (H, I). *n* = 18 embryos per staining. **(H)** Schematic illustrating the strategy used for *ptc2* quantification, with regions of interest (ROIs) drawn over EGFP^+^ cells and immediately adjacent EGFP^-^ control regions. **(I)** Paired comparison of *ptc2* fluorescence intensity in EGFP^+^ versus EGFP^-^ ROIs, showing reduced Shh pathway activity in Foxa2-overexpressing cells. Each data point represents one paired ROI measurement. *n* = 37 paired measurements from 18 embryos. Data in (C, E, F, I) are plotted as mean ± SD. Statistics: Mann–Whitney U test (C, E, and paired t-test (I). Significance: p > 0.05 (ns), p < 0.0001 (****). Scale bars: 20 μm (A, B,; 100 μm (D).

Since *foxa2*-deficient embryos show increased Shh signaling in the ventral spinal cord, we asked whether Foxa2 overexpression alters Shh signaling responsiveness at the cellular level. To address this, we quantified Shh pathway activity using *ptc2* expression as a readout in embryos mosaically expressing *hsp:foxa2-P2A-EGFP*. Specifically, we measured *ptc2* fluorescence intensity within circular regions centered on EGFP-positive *foxa2^GOF^*cells and compared these values to adjacent EGFP-negative regions (Fig. 6G and 6H). Paired t-test analysis revealed a significant reduction in *ptc2* expression in EGFP-positive regions (Fig. 6I), suggesting that Foxa2 negatively regulates Shh responsiveness in the ventral spinal cord cell-autonomously.

Collectively, these gain-of-function results are consistent with our loss-of-function analyses and support a model in which Foxa2 is sufficient to promote floor plate identity, drive KA” interneuron differentiation, and attenuate Shh pathway responsiveness in a cell-autonomous manner.

## DISCUSSION

Previous studies in zebrafish and other vertebrates have established several foundational aspects of Foxa2 function in midline development, including its expression in the FP and its requirement for FP marker expression^25,44,60,61^. These studies are essential for defining Foxa2 as a conserved regulator of axial mesoderm and FP development^25,31,33,44,45^. However, several mechanistic questions remain unresolved. It is unclear whether prospective FP cells are lost, mispositioned, or transformed in the absence of Foxa2. It is also unknown whether Foxa2 performs distinct functions in the neighboring neural p3 domain. Here, we demonstrate that Foxa2 acts not only as a regulator of FP marker expression but also as a lineage barrier that prevents mesoderm-derived FP cells from acquiring Shh-responsive neural progenitor identities. In addition, we identify a separate role for Foxa2 in the p3 lineage, where it attenuates Shh signaling responsiveness to promote differentiation of KA” interneurons. Thus, our work extends previous studies by showing that loss of Foxa2 does not simply abolish FP marker expression; rather, it reveals a broader role for Foxa2 in maintaining lineage identity, limiting Shh responsiveness, and promoting context-dependent neural differentiation.

### Foxa2 preserves FP identity by preventing neural fate transformation

Previous studies have shown that loss of Foxa2 disrupts the expression of canonical FP markers, including *shh*-family genes, in zebrafish and other vertebrates^25,33,44,45^. Our results confirm this conserved requirement while further clarifying the cellular basis of the phenotype. Rather than indicating that prospective FP cells fail to form, die, or migrate incorrectly, our lineage-tracing experiments show that shield-derived cells still reach the ventral midline in Foxa2-deficient embryos. However, these cells fail to maintain FP identity and instead acquire features of neural progenitors and neurons. This distinction is important because it indicates that Foxa2 is required not only for the FP differentiation program but also for suppressing alternative neuroectodermal fates in mesoderm-derived midline cells. Indeed, depending on Shh signaling levels, these transformed cells can adopt p3, pMN, or even more dorsal identities. The ability of FP cells to convert into multiple ventral and dorsal neural identities upon loss of Foxa2 reveals an unexpected flexibility within the ventral midline. Our findings thus provide a mechanistic explanation for the long-observed loss of FP markers in *foxa2* mutants.

Similar transcription factor-dependent fate switches have been described in other systems. In the adult Drosophila midgut, absence of transcriptional repressor Ttk69 redirects intestinal stem cell lineages toward enteroendocrine cell fates, demonstrating that loss of a single factor can reprogram lineage output^62,63^. A comparable example is observed in the mouse retina, where a transcription factor called Fezf1 controls a postmitotic binary fate choice between closely related interneuron subtypes^64^. Loss of Fezf1 causes neurons that normally adopt one interneuron identity to switch to an alternative subtype^64^. Notably, these examples involve fate transitions between closely related cell types within the same lineage, whereas our findings reveal a broader plasticity in which mesoderm-derived floor plate cells adopt neuroectodermal identities.

### Sequential roles of Foxa2 in regulating neural progenitor differentiation

In contrast to its fate-inducing role in the FP, Foxa2 performs a distinct function within the adjacent p3 neural progenitor domain. Loss of *foxa2* resulted in elevated Shh pathway activity in the ventral spinal cord, despite the concomitant loss of the FP, a major source of Shh ligand. This finding implicates Foxa2 as a negative regulator of Shh responsiveness. Consistent with this, ectopic Foxa2 expression led to reduced expression of the Shh target *ptc2* in a cell-autonomous manner, indicating that Foxa2 is sufficient to suppress Shh pathway activity. Our results reveal that Foxa2 both positively and negatively regulates the Shh pathway, depending on cellular context. In the FP, Foxa2 promotes *shh* ligand expression, as shown by our data and supported by a previous *foxa2* misexpression study^61^. At the same time, Foxa2 suppresses Shh signaling response within both FP and p3 cells, preventing FP to acquire a neural fate and promoting p3 cells to differentiate into neurons. This dual role is consistent with findings from the mouse ventral midbrain, where Foxa1 and Foxa2 are shown to activate Shh expression^65^. They also repress Shh signaling responses through direct regulation of intracellular pathway components such as Gli2^65^. Similarly, studies in the chick spinal cord have shown that ectopic Foxa2 expression leads to downregulation of the Shh target *Ptch1*, whereas expression of a dominant-negative form of Foxa2 results in ectopic *Ptch1* expression, indicating that Foxa2 can negatively regulate Shh pathway output downstream of ligand production^66^. Together, these findings demonstrate that regulation of Shh signaling by Foxa2 is a conserved feature of vertebrate neural development.

The p3 domain further differentiates into KA” interneurons, which are considered the zebrafish counterparts of cerebrospinal fluid-contacting neurons (CSF-cNs) in mammals^16,17,20,67^. Interestingly, Foxa2 expression is specifically upregulated in KA” interneurons compared with p3 progenitors. Our previous work has shown that neural differentiation requires timely attenuation of the Shh response^16,50^. The elevated Foxa2 expression in KA” cells suggest a model where Foxa2 promotes the differentiation of p3 progenitors to KA” interneurons by inhibiting Shh pathway activity. Indeed, despite having more p3 progenitors, *foxa2* mutants show a reduction in the number of KA” interneurons. Intriguingly, in Foxa2-deficient embryos, KA” interneurons are preferentially positioned at the ventral midline rather than forming two lateral rows as observed in controls. The mechanisms underlying this altered spatial organization are currently unknown, but we speculate that the absence of the FP barrier at the midline, as well as Notch-mediated lateral inhibition, may contribute to this phenotype.

Because ectopic Foxa2 overexpression can induce the KA”-specific neuropeptide genes *urp1* and *urp2*^68^ in a stage- and tissue-independent manner, this suggests that Foxa2 may directly activate transcriptional programs required for KA” function. Therefore, our results support that Foxa2 coordinates p3 differentiation through a two-step mechanism: first by attenuating Shh signaling to permit neural differentiation, and subsequently by activating KA”-specific transcriptional programs that reinforce the neural identity. Importantly, this function is not restricted to the vertebrate spinal cord. In *C. elegans*, the FoxA homolog PHA-4, best known for its role in early gut patterning, also functions in the postmitotic enteric neurons^69^. PHA-4 initiates terminal differentiation of pharyngeal enteric neurons and maintains neuronal function in GABAergic hindgut neurons to regulate defecation behavior^69^. Together, these observations support a conserved role for FoxA transcription factors in promoting neuronal differentiation and maintaining neural identity across species.

### Temporal and context-dependent functions of Foxa2 as a pioneer transcription factor

An important question raised by our findings is why Foxa2 expression in the p3 domain does not convert p3 progenitors into FP cells, despite Foxa2 being sufficient to induce FP fate when ectopically expressed. Our data suggest that developmental timing and cellular context are the critical determinants. In the FP, *foxa2* expression begins early, at the shield stage, under the control of Nodal signaling^21,24,60,70,71^. In contrast, *foxa2* expression in the p3 domain is Shh-dependent and occurs later^21,72^. Consistent with this temporal regulation, our gain-of-function experiments demonstrate that only ‘early’ ectopic expression of Foxa2 is sufficient to induce FP marker expression. Late expression, corresponding to the timing of endogenous p3 Foxa2 expression, is insufficient. These findings suggest that cells possess a transient competence to adopt FP identity that is progressively restricted as neural progenitor programs become established. Once this competence is lost, Foxa2 expression alone is no longer sufficient to redirect cell fate.

More broadly, Foxa2’s ability to perform distinct functions in adjacent domains is consistent with its role as a pioneer transcription factor capable of shaping regulatory landscapes. We propose that early Foxa2 activity establishes a FP-specific chromatin state before neural progenitor identities are consolidated, thereby stabilizing midline fate. In contrast, later Foxa2 expression in p3 progenitors may operate within a chromatin environment already configured for neural differentiation. Supporting this model, Delás *et al.* demonstrate that FOXA2 opens p3-specific regulatory elements that subsequently recruit NKX2.2, establishing the p3 transcriptional program in embryonic stem cell-derived neural progenitors^23^. A similar context-dependent regulatory logic has been described in lung adenocarcinoma, where FOXA1/2 cooperate with the pulmonary lineage factor NKX2-1 to maintain lung tumor identity, but in the absence of NKX2-1, FOXA1/2 drive epigenomic reprogramming and activate a gastric-like lineage program^73,74^. Together, these findings support a model in which Foxa2 confers lineage competence rather than instructing a fixed identity, allowing cells to integrate temporal and positional cues with intrinsic transcriptional states. Although we did not directly assess chromatin accessibility, our findings are consistent with a model in which temporal differences in Foxa2 activity define windows of cellular competence. An important next step for us will be to identify the unique transcriptional targets of Foxa2 in the FP and p3 domains to understand how context and timing influence its function.

In summary, our work identifies Foxa2 as a dual-function regulator of ventral spinal cord development. Foxa2 maintains floor plate identity and prevents Shh-dependent neural differentiation, while simultaneously promoting interneuron differentiation in the adjacent p3 domain through modulation of Shh signaling responsiveness. These findings illustrate how a single transcription factor can generate distinct developmental outcomes in neighboring cell populations through temporally restricted windows of cellular competence.

## MATERIALS AND METHODS

### Ethics statement

All animal research was conducted in accordance with the principles outlined in the current Guideline of the Canadian Council on Animal Care. All protocols were approved by the Animal Care Committee at the University of Calgary (AC21-0102 and AC25-0038).

### Zebrafish strains

Zebrafish strains used in this research were raised under standard conditions^75^. The following transgenic strains were used in our study: *TgBAC(ccn2a:EGFP)ca119, Tg(gata2:GFP)la3*^76^, *Tg(nkx2.2a:NLS-mCherry)ca114*^18^, *Tg(phiC31.attP1)co2002*^77^ (referred to as *pIGLET24b*), *TgBAC(ptc2:Kaede)a4596*^16^, and *Tg(slc17a6b:EGFP)zf139*^78^ (referred to as *vglut2a:GFP*). The *foxa2^st^*^20^ mutant^44,79^ and *igu^ts^*^294^ mutant^54,55^ strains were maintained as heterozygotes, and homozygous embryos (referred to as *foxa2^-/-^*and *igu^-/-^*) were generated by intercrossing heterozygous carriers.

### Mutant genotyping

Genomic DNA was extracted from embryos or adult fin clips using the HOTSHOT protocol. Briefly, tissue was incubated in 50 mM NaOH at 95 °C for 20 min, followed by cooling on ice and neutralization with 1 M Tris-HCl (pH 7.5). PCR amplification was performed using specific primer pairs, followed by restriction enzyme digestion.

*foxa2^st^*^20^ genotyping: The *foxa2^st^*^20^ allele contains a single C-to-A point mutation that disrupts the *foxa2* gene^44,79^. A region flanking the mutation site was amplified using the forward primer (5′-CATGAACACTTACATGACTATGTCCG-3′) and reverse primer (5′-AGCGTTGCTGGTTCTGTCG-3′). PCR reactions were performed with an annealing temperature of 58 °C. The PCR product was digested with MseI at 37 °C. The wild-type allele produced a single 464 bp band, whereas the mutant allele generated two fragments of 412 bp and 52 bp. Heterozygous embryos displayed all corresponding bands.

*igu^ts^*^294^ genotyping: The *igu^ts^*^294^ allele carries a single point mutation introducing a premature stop codon in the *igu* gene^54,55^. A region spanning the mutation site was amplified using the forward primer (5′-TGCTGAAACTGCAGCAGAAA-3′) and reverse primer (5′-TGGATGAAAAGCATCGTCAA-3′), with an annealing temperature of 58 °C. The PCR product was digested with BsaXI at 37 °C. The wild-type allele produced three fragments of 285 bp, 97 bp, and 30 bp, and the mutant allele remained undigested at 412 bp. Heterozygous embryos displayed all corresponding bands.

### Generation of transgenic lines

To generate the *ccn2a:EGFP* transgenic line, the BAC clone zK266K12 from the Danio Key library, which contains the *ccn2a* genomic region, was selected for bacteria-mediated homologous recombination following standard protocols^80^. First, a cassette containing two oppositely oriented Tol2 arms and an ampicillin resistance gene was recombined into the BAC vector. Then, a cassette containing EGFP and a kanamycin resistance gene was recombined into the BAC to replace the first coding exon of the *ccn2a* gene. Successfully modified BAC constructs were confirmed by PCR analysis. The recombinant *ccn2a:EGFP* BAC was then co-injected with *tol2* transposase mRNA into wild-type embryos at the one-cell stage. Stable transgenic lines were established by screening F1 embryos from injected founders for EGFP expression.

### Morpholino and plasmid injections

To knock down *foxa2*, morpholino oligonucleotides (Gene Tools, LLC) targeting *foxa2* (5’-CCTCCATTTTGACAGCACCGAGCAT-3’)^61^ were injected at 0.8 mM into one-cell stage embryos with 1 nL per embryo. Injected embryos were fixed at appropriate stages for in situ hybridization analysis.

For overexpression experiments, a *hsp:foxa2-P2A-EGFP-attB* plasmid was generated using Gibson assembly (NEB #E2611). 40 pg of *hsp:foxa2-P2A-EGFP-attB* plasmid DNA was co-injected with 25 pg of *phiC31* integrase mRNA into *pIGLET24b* embryos carrying the attP landing site at the one-cell stage. Injected embryos were heat shocked at appropriate stages for analysis.

### In situ hybridization and immunohistochemistry

Whole-mount in situ hybridization were performed following standard protocols^81^. The RNA probes used were: *foxa2*, *irx3*, *nkx2.2a*, *nkx2.2b*, *olig2*, *pax6*, *ptc2*, *shha*, *shhb*, *sim1*, *spon1b*, *urp1*, and *urp2.* Double fluorescent in situ hybridizations were performed by combining different digoxigenin (DIG)- and dinitrophenyl (DNP)-labeled probes with homemade FITC and Cy3 tyramide solutions^82^. For immunohistochemistry, the primary antibodies used were: rabbit monoclonal anti-Foxa2 (1:100, Abcam, ab108422), chick polyclonal anti-GFP (1:250, Aves, GFP-1020), rabbit polyclonal anti-GABA (1:500, Sigma, A2052), mouse monoclonal anti-CaMPARI-red (1:100, gift from Eric Schreiter), mouse monoclonal anti-HuC/D (1:100, Invitrogen, A21271), and rabbit polyclonal anti-fluorescein (1:50, Invitrogen, A-889). Appropriate Alexa Fluor-conjugated secondary antibodies (1:500, Thermo Fisher) were used for fluorescent detection of antibody labeling. Draq5 (1:5000, Biostatus, DR50050) was used for nuclear staining.

### Embryo sectioning

To obtain transverse views, stained embryos were manually sectioned using vibratome steel blades or a cryostat. For cryosectioning, embryos were cryoprotected in 30% sucrose at 4 °C before being embedded in OCT compound (VWR) and frozen at −80 °C. Sections (10-16 µm) were cut using a Leica cryostat. Sections were collected from the trunk region dorsal to the yolk extension.

### Fate-mapping experiments

Fate-mapping experiments were performed using caged fluorescein dextran photoactivation and Kaede photoconversion. For uncaging experiments, caged fluorescein dextran was generated by conjugating CMNB-caged fluorescein to aminodextran following a previously described protocol^83^. Embryos were injected at the one-cell stage with caged fluorescein dextran (1:2 dilution) and maintained at 28.5 °C in the dark. At the shield stage, embryos were mounted dorsal side down in agarose wells generated using 3D-printed molds (gift from Joshua Bloomekatz) on a glass-bottomed dish. A defined ∼60-µm region of interest (ROI) within the shield, corresponding to approximately 8-12 cells, was selected for photoactivation. Illumination was performed using a 405-nm UV laser on an Olympus FV1200 inverted confocal microscope. Successful photoactivation was confirmed immediately after illumination, and embryos were returned to 28.5 °C for further development. Labeled cells were followed over time, and cell fate was assessed at the appropriate stages by live imaging or by immunohistochemistry.

For fate-mapping using Kaede photoconversion, Kaede mRNA was synthesized in vitro using the mMESSAGE mMACHINE kit (Ambion). A total of 100-120 pg of Kaede mRNA were injected at the one-cell stage. Photoconversion experiments were carried out similarly as described above.

### Drug treatment

To inhibit Shh signaling, embryos at 6 hpf were treated with cyclopamine (Toronto Chemical, 100 μM) or DMSO (control) in E3 fish water until 27 hpf and fixed at 28 hpf for analysis.

### Heat shock induction

To induce expression from the heat shock promoter, *hsp:foxa2-P2A-EGFP*-injected embryos at the relevant stage were placed in a 2 mL micro-centrifuge tube in a heat block set to 37 °C for 30 min. After heat shock, embryos were transferred back into E3 fish water in a Petri dish and allowed to recover at 28.5 °C. *hsp:foxa2-P2A-EGFP*-positive embryos were selected based on EGFP expression 3 hours after heat shock.

### Quantification of *ptc2:Kaede* levels

To quantify *ptc2:Kaede* expression across the dorsoventral axis of the spinal cord, a modified version of Cimtool (MATLAB-based) was used^59^. Multipage TIFF files were separated into two image stacks: a brightfield channel and a gene expression channel.

For spinal cord segmentation, a z-projection of the brightfield channel was generated to obtain a single 2D image. The projected image was filtered using a 3 × 3 median filter to reduce noise while preserving edges^58^. Points along the bottom edge of the spinal cord were manually selected and used, together with the filtered image, as inputs for the contour segmentation algorithm implemented in Cimtool. The upper edge was segmented using the same procedure.

For gene expression analysis, the start and end positions of the image stack were defined to include the region with the highest *ptc2:Kaede* expression. To minimize overlap between adjacent optical sections, every fifth slice within this range was analyzed. Within each selected image, a region of interest corresponding to the area of highest reporter expression was defined. Line segments perpendicular to the bottom edge were then generated from the bottom to the top boundary of the spinal cord. Gene expression intensity profiles along these segments were extracted using the MATLAB function *improfile*.

For each image, the extracted profiles and associated descriptors (maximum intensity, profile length, and magnitude and position of the negative gradient) were exported as CSV files^84^. For each embryo, intensity profiles from all analyzed images were averaged to generate a mean dorsoventral expression profile. Group-level expression profiles were calculated by averaging the mean profiles across embryos for each condition (control and *foxa2^MO^*). In addition, the median values of the maximum intensity and negative gradient descriptors were calculated for each embryo and plotted for comparisons.

The modified Cimtool MATLAB source code is available at: https://github.com/lucaslovercio/SpinalCordAnalysis

### Quantification of KA” and V3 interneurons

To quantify KA” and V3 interneurons, embryos were stained with appropriate markers (KA”: *urp1*; V3: *sim1*). Labeled neurons were manually counted within the spinal cord region, from the anterior-most labeled neuron to the end of the yolk extension for each embryo.

### Quantification of p3 cells

To quantify p3 cells, embryos were stained for *nkx2.2a* and the nuclear marker Draq5. Counts were performed within a defined three-somite region per embryo. Nuclei were identified by Draq5 staining, and only Draq5^+^ nuclei co-expressing *nkx2.2a* were included. Nuclei were counted using the cell counter tool in Fiji/ImageJ^85^. Counts from individual embryos were used for statistical analysis.

### Quantification of marker expression in *foxa2^GOF^* embryos

To assess the induction of FP and KA”-associated markers following Foxa2 overexpression, multiple complementary quantification approaches were used as described below.

Induction of FP and KA” markers: To quantify induction of FP and KA”-associated markers following Foxa2 overexpression, the proportion of EGFP-positive cells expressing *shha*, *shhb, spon1b*, or *urp1* was determined from confocal images. Quantification was restricted to the spinal cord region of each embryo. For each image, the EGFP channel was examined first, and all EGFP-positive cells were manually identified and labeled using the cell counter tool in Fiji/ImageJ^85^. EGFP-positive cells were scored as marker-positive if overlapping signal was detected in the corresponding channel. The percentage of EGFP-positive cells expressing each marker was calculated for each embryo and plotted in the graph.

Quantification of ectopic *shha* expression: Ectopic *shha* expression was quantified from whole-mount NBT/BCIP in situ hybridization images by manual counting. For each embryo, discrete *shha^+^* clusters located outside the endogenous floor plate and notochord expression domains were counted and recorded as the number of ectopic *shha^+^* clusters per embryo.

Quantification of ectopic *urp1* expression: Ectopic *urp1* expression was quantified from whole-mount NBT/BCIP in situ hybridization images using image-based threshold analysis in Fiji/ImageJ^85^. Images were duplicated and converted to 8-bit grayscale, and an intensity threshold was applied to capture *urp1* staining while minimizing background and yolk-associated signal. A polygonal ROI was manually drawn to encompass the embryo body while excluding the yolk, and this ROI was applied to the thresholded binary image for measurement. This ensured that yolk-associated signal was excluded from all analyses. For each time point, the average *urp1* staining area measured in the control embryos was calculated and subtracted from individual measurements obtained from *foxa2^GOF^* embryos to remove endogenous *urp1* expression and isolate ectopic staining. Data are presented as the percentage of ectopic *urp1^+^* staining area per embryo.

Quantification of *ptc2* expression in mosaic Foxa2-overexpressing embryos: *ptc2* expression was quantified from confocal images of embryos mosaically expressing *hsp:foxa2-P2A-EGFP*. A single optical section was used for each measurement. Circular ROIs were manually drawn around individual EGFP^+^ cell clusters, and mean *ptc2* fluorescence intensity was measured within each ROI. For each EGFP^+^ ROI, two immediately adjacent EGFP^-^ ROIs of identical size were placed in the same optical section, and the mean of these two measurements was used as the local control. ROI size was kept identical between EGFP^+^ and adjacent EGFP^-^ regions for each paired measurement. A total of 37 paired measurements from 18 embryos were analyzed. Statistical significance was assessed using a two-tailed paired t-test (GraphPad Prism).

### Statistical analysis

All the graphs were generated in the GraphPad Prism software. Data were plotted as mean ± SD. Significance between two samples was calculated using Mann-Whitney U test. A two-tailed paired t-test was used for Fig. 6I. p values: p > 0.05 (not significant, ns); p < 0.05 (∗); p < 0.01 (∗∗); p < 0.001 (∗∗∗); p < 0.0001 (∗∗∗∗).

## ACKNOWLEDGMENTS

We thank the zebrafish community for sharing reagents, particularly the Zebrafish International Resource Center (ZIRC) for the *foxa2* mutant and Christian Mosimann for the *pIGLET24b* line. We are grateful to Eric Schreiter for generously providing the anti-CaMPARI-red antibody, to Joshua Bloomekatz for sharing the 3D-printed molds used for embryo mounting, and to Tika Kocha for excellent zebrafish care. We also thank Sarah Childs for critical input on this project, as well as members of the Childs and Huang laboratories for valuable discussions.

---

**Figure S1.**
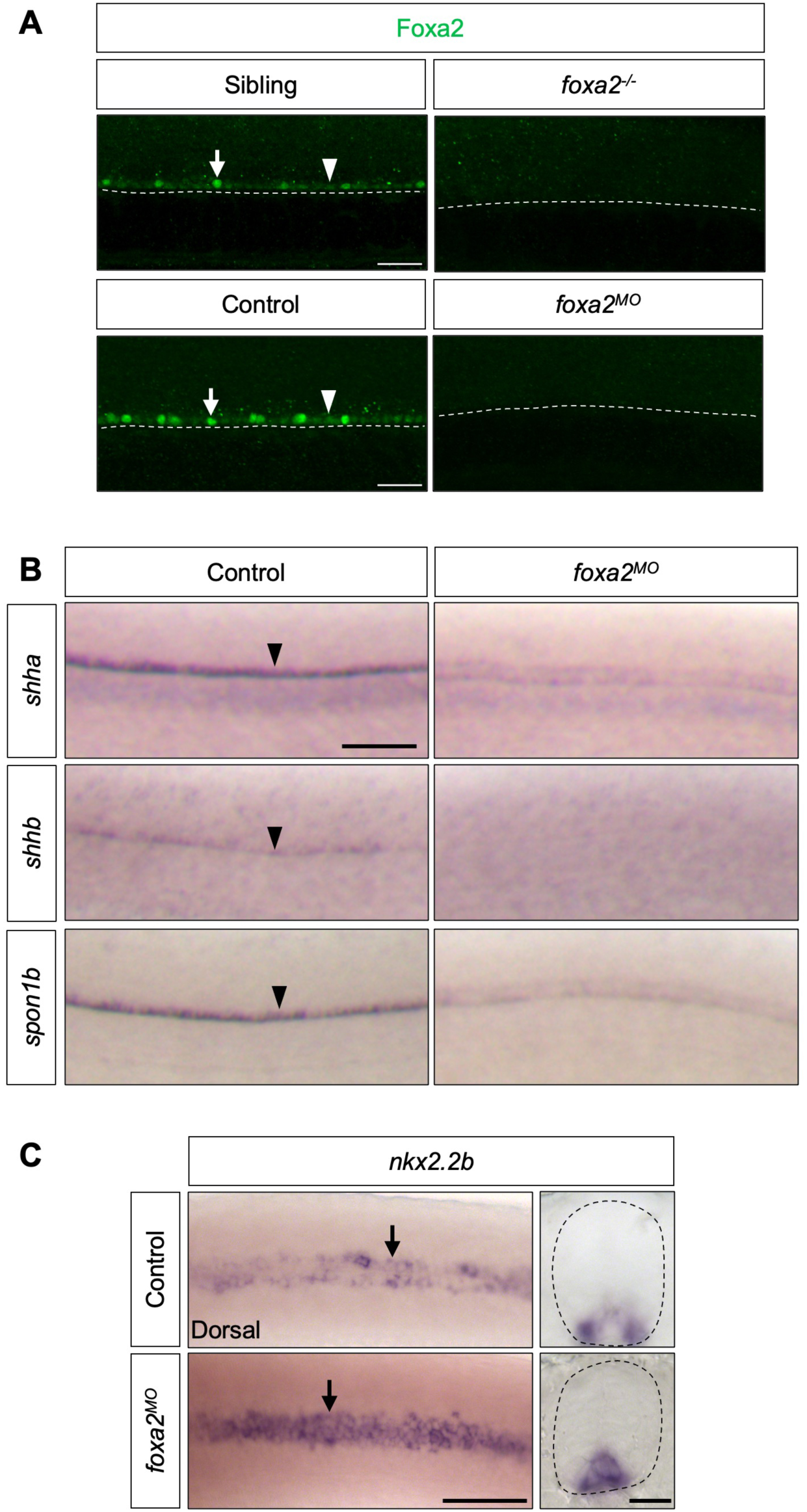
*foxa2* morpholino knockdown phenocopies *foxa2* mutants and disrupts FP and p3 patterning. **(A)** Representative lateral views of Foxa2 immunostaining at 30 hpf show loss of Foxa2 protein in both *foxa2^-/-^* mutants and *foxa2^MO^* embryos. Arrowheads indicate Foxa2 expression in the FP, whereas arrows indicate KA” cells with higher Foxa2 expression. Dashed lines mark the boundary between the FP and notochord. *n* = 10 embryos per staining. **(B)** Whole-mount in situ hybridization for the FP markers *shha*, *shhb*, and *spon1b* in control and *foxa2^MO^*-injected embryos at 30 hpf, showing loss or reduction of FP marker expression (arrowheads) following *foxa2* knockdown. Representative lateral views are shown. *n* = 20 embryos per staining. **(C)** Whole-mount in situ hybridization for the p3 marker *nkx2.2b* in control and *foxa2^MO^*embryos. Dorsal views (left) and transverse sections (right) show expansion of *nkx2.2b* expression (arrows) into the ventral midline following Foxa2 loss. Dashed outlines indicate the spinal cord. *n =* 20 embryos per condition. Scale bars: 50 μm (A); 100 μm (B); 50 μm (dorsal view) and 20 μm (transverse view) (C).

**Figure S2.**
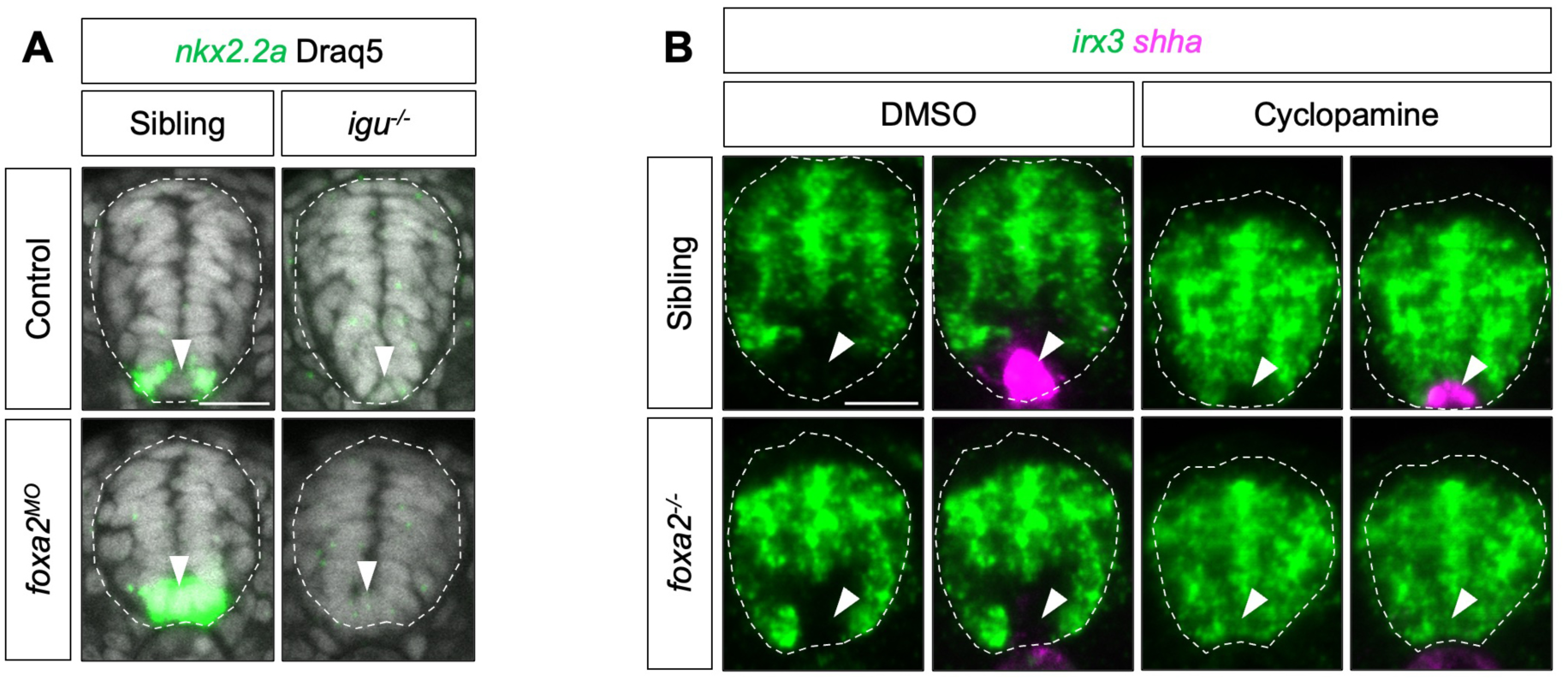
Fate-switched FP cells acquire Shh signaling responsiveness in the absence of Foxa2. **(A)** Representative transverse sections showing *nkx2.2a* expression in uninjected and *foxa2^MO^*-injected sibling controls (*igu^+/+^* or *igu^+/-^*) and homozygous *igu^-/-^* mutants at 28 hpf. Nuclei are marked by Draq5 staining. Arrowheads indicate the ventral midline of the spinal cord. *n* = 5 embryos per condition. **(B)** Fluorescent in situ hybridization for dorsal spinal cord marker *irx3* (green) and the FP marker *shha* (magenta) in sibling control and *foxa2^-/-^*embryos treated with DMSO or cyclopamine from 6 hpf to 27 hpf and fixed at 28 hpf. *n* = 5 embryos per condition. Dashed outlines indicate the spinal cord, and arrowheads point to the ventral midline of the spinal cord. Scale bars: 20 μm.

**Figure S3.**
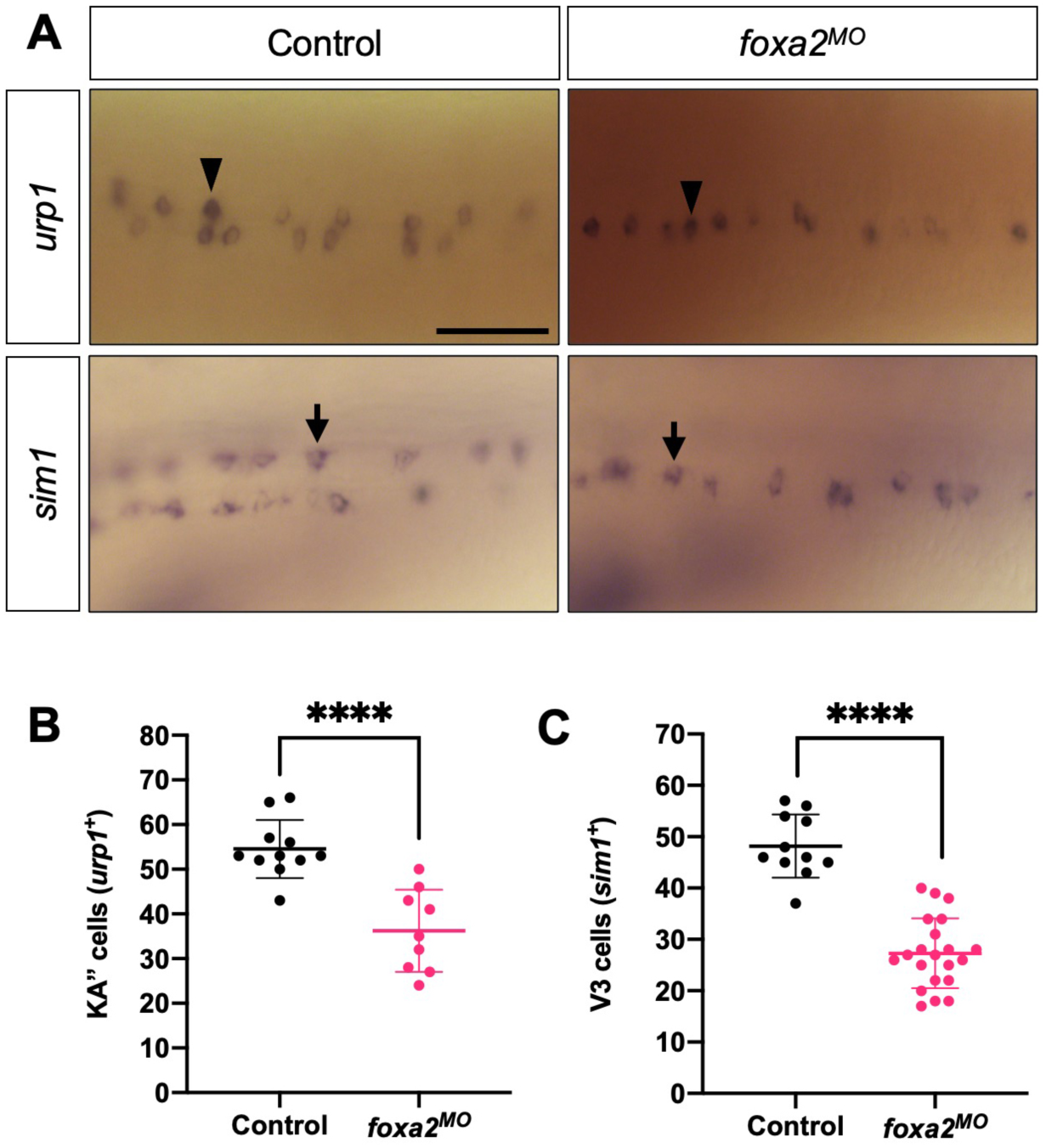
*foxa2* knockdown results in fewer KA” and V3 interneurons. **(A)** Whole-mount in situ hybridization for the KA” interneuron marker *urp1* (arrowheads) and the V3 interneuron marker *sim1* (arrows) in control and *foxa2^MO^* embryos at 30 hpf. Dorsal views of the spinal cord are shown. Quantification of KA” and V3 cell numbers is shown in (B, C). *n* = 25 embryos per condition. **(B)** Quantification of *urp1^+^*KA” interneuron numbers. Each data point represents one embryo. *n* = 11 (control) and 9 (*foxa2^MO^*) embryos. **(C)** Quantification of *sim1^+^* V3 interneuron numbers. Each data point represents one embryo. *n* = 11 (control) and 21 (*foxa2^MO^*) embryos. Data are plotted as mean ± SD. Statistics: Mann–Whitney U test. Significance: p < 0.0001 (****). Scale bar: 50 μm.

**Figure S4.**
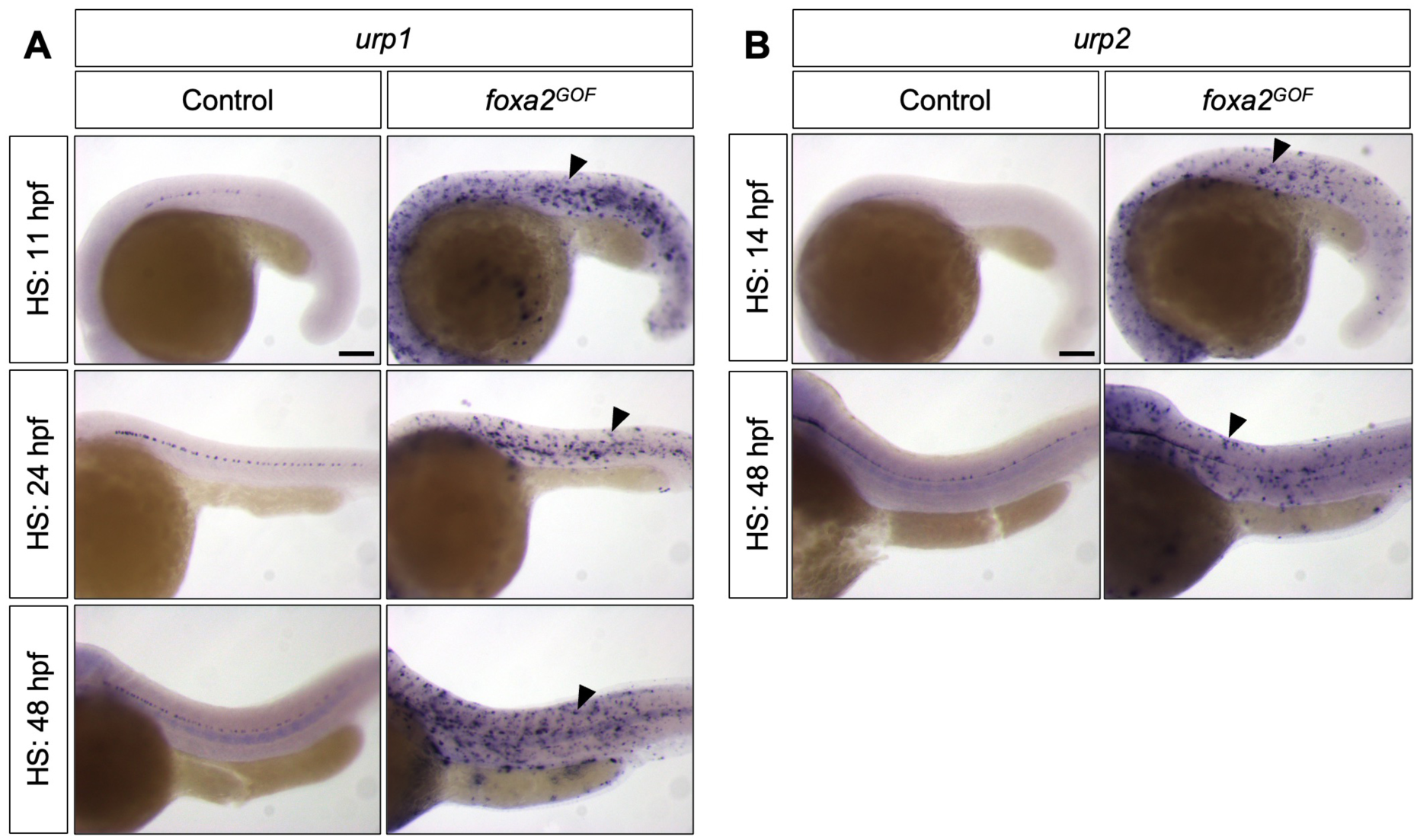
Foxa2 overexpression induces widespread ectopic KA” marker expression. **(A)** Whole-mount in situ hybridization for the KA” marker *urp1* in control and *hsp:foxa2-P2A-EGFP*-injected (*foxa2^GOF^*) embryos following heat shock at the indicated developmental stages (11, 24, and 48 hpf) and fixation 6 hours later, revealing robust ectopic *urp1* expression (arrowheads) throughout the embryo. *n* ≥ 12 per staining. **(B)** Whole-mount in situ hybridization for the KA” marker *urp2* in control and *foxa2^GOF^* embryos following heat shock at the indicated developmental stages (14 and 48 hpf) and fixation 6 hours later, showing robust ectopic *urp2* expression (arrowheads) throughout the embryo. *n* ≥ 10 per staining. Scale bars: 100 μm.

